# APOE genotype-dependent differences in human astrocytic energy metabolism

**DOI:** 10.1101/2025.03.31.646283

**Authors:** Vanessa Budny, Chantal Bodenmann, Kathrin J. Zürcher, Sherida M. de Leeuw, Rebecca Z. Weber, Luca Ravotto, Iván Ruminot, L. Felipe Barros, Bruno Weber, Christian Tackenberg

**Affiliations:** Institute for Regenerative Medicine, University of Zurich, 8952 Schlieren, Switzerland; Neuroscience Center Zurich, University of Zurich and ETH Zurich, Zurich, Switzerland; Institute for Pharmacology and Toxicology, University of Zurich, 8057 Zürich, Switzerland; Centro de Estudios Científicos (CECs), Valdivia, Chile; Facultad de Ciencias para el Cuidado de la Salud, Universidad San Sebastián, Valdivia, Chile; Facultad de Medicina, Universidad San Sebastián, Valdivia, Chile

**Keywords:** Apolipoprotein E (APOE), Alzheimer’s disease (AD), induced pluripotent stem cells (iPSCs), human astrocytes, energy metabolism, glycolysis, mitochondrial function, fluorescence lifetime imaging (FLIM), Seahorse assay

## Abstract

The main genetic risk factor for Alzheimer’s disease (AD) is the presence of the apolipoprotein E4 (*APOE4*) allele. While *APOE4* increases the risk of developing AD, the *APOE2* allele is protective and *APOE3* is risk-neutral. In the brain, APOE is primarily expressed by astrocytes and plays a key role in various processes including cholesterol and lipid transport, neuronal growth, synaptic plasticity, immune response and energy metabolism. Disruptions in brain energy metabolism are considered a major contributor to AD pathophysiology, raising a key question about how different APOE isoforms affect the energy metabolism of human astrocytes.

In this study, we generated astrocytes (iAstrocytes) from *APOE*-isogenic human induced pluripotent stem cells (iPSCs), expressing either APOE2, APOE3, APOE4 or carrying an APOE knockout (*APOE-KO*), and investigated *APOE* genotype-dependent changes in energy metabolism. ATP Seahorse assay revealed a reduced mitochondrial and glycolytic ATP production in *APOE4* iAstrocytes. In contrast, proteomic GO enrichment analysis and mitochondrial stress tests indicated increased mitochondrial respiration and activity in *APOE4* iAstrocytes, accompanied with elevated proton leak, while mitochondrial fusion and fission protein levels remain unchanged. Glycolysis stress tests also demonstrated enhanced glycolysis and glycolytic capacity in *APOE4* iAstrocytes while genetically encoded nanosensor-based FLIM analysis revealed that *APOE* does not affect lactate dynamics. Mass spectrometry-based metabolomic analysis identified various energy and glucose metabolism-related pathways that were differentially regulated in *APOE4* compared to the other genotypes, including mitochondrial electron transport chain and glycolysis. In general, *APOE2* and *APOE-KO* iAstrocytes showed a very similar phenotype in all functional assays and differences between *APOE2*/*APOE-KO* and *APOE4* were stronger than between *APOE3* and *APOE4*.

Our study provides evidence for *APOE* genotype-dependent effects on astrocyte energy metabolism and highlights alterations in the bioenergetic processes of the brain as important pathomechanisms in AD.

## 1. Introduction

The human brain is a highly energy-demanding organ and requires substantial metabolic resources to maintain its functions. Alterations in the bioenergetic processes of the brain are commonly observed during aging and are implicated in various neurodegenerative diseases, such as Alzheimer’s disease (AD). AD is the most common age-related neurodegenerative disease affecting memory and executive functions. Several genetic and non-genetic risk factors have been identified so far to influence the AD risk. Among those, the ε4 allele of the apolipoprotein E (*APOE4*) represents the strongest genetic risk factor (Tanzi, 2012). Three major allelic variants of human *APOE* have been identified: *APOE2*, *APOE3* and *APOE4*. *APOE4* increases the risk for developing AD by 2.5-4-fold in heterozygous and 12-15-fold in homozygous carriers (Corder et al., 1993; Grupe et al., 2007; Corneveaux et al., 2010), however the risk is affected by factors such as sex or population ancestry (Belloy et al., 2023). In contrast, *APOE3* is defined as risk-neutral while *APOE2* represents the strongest genetic protective factor (Corder et al., 1994; Farrer et al., 1997; Reiman et al., 2020). In the brain, APOE is mainly produced by astrocytes, which generate up to 80% of total brain APOE (Blumenfeld et al., 2024). APOE isoforms differentially affect astrocyte function. Human isogenic iPSC-derived astrocytes expressing APOE4 displayed a pro-inflammatory phenotype, reduced beta-amyloid update capacity and altered lipid metabolism compared to astrocytes expressing APOE3 or APOE2 (de Leeuw et al., 2022). Importantly, *APOE* is also implicated in regulating bioenergetic homeostasis of the brain.

Numerous studies have demonstrated altered glucose metabolism as a common phenotype in AD, especially in *APOE4* carriers. [^18^F]fluorodeoxyglucose (FDG)-PET imaging showed hypometabolism in AD brains, mainly in the temporoparietal cortex including the precuneus and posterior cingulate cortex (Reiman et al., 1996; Foster et al., 2007). Interestingly, several studies in young *APOE4* carriers or in *APOE4*-knock-in mice with no apparent neurodegenerative pathology, revealed increased brain metabolism and activity (Filippini et al., 2009; Evans et al., 2017; Nuriel et al., 2017; Venzi et al., 2017), indicating that the APOE4 effect on brain energy metabolism may differ depending on the degree of pathology. Further, post-mortem analysis of AD brains discovered higher brain tissue glucose concentration, reduced glycolytic flux and lower levels of neuronal glucose transporter GLUT3 but no effect on astrocytic glucose transporter GLUT1, suggesting cell type specific metabolic alterations in AD (An et al., 2018).

The proposed mechanisms of how *APOE* variants, especially *APOE4*, affect the energy metabolism of neural cells remain inconclusive. They range from reduced glycolysis in APOE4-expressing cells (Wu et al., 2018; Fang et al., 2021; Zhang et al., 2023), via a metabolic shift from oxidative phosphorylation to glycolysis in *APOE4* or AD patient cells (Sonntag et al., 2017; Williams et al., 2020; Farmer et al., 2021; Lee et al., 2023) to an increase in oxidative phosphorylation in *APOE4* induced neurons (Budny et al., 2024). Further, many studies only compared two *APOE* variants, mostly *APOE4* and *APOE3*. Thus, more research is required to specifically show how *APOE4*, compared to both *APOE3* and *APOE2*, alters glycolytic and mitochondrial function in human brain cells.

In the present study, we used human APOE isogenic iPSC-derived astrocytes (iAstrocytes) to investigate the impact of *APOE4*, *APOE3*, *APOE2* and *APOE-KO* on astrocyte energy metabolism. Our findings reveal *APOE* genotype-dependent metabolic and proteomic changes. *APOE4* iAstrocytes showed upregulated oxidative phosphorylation (OXPHOS) proteomic pathways, accompanied by increased mitochondrial respiration and glycolysis. Interestingly, despite the elevated metabolic activity, *APOE4* iAstrocytes produced less ATP compared to the other *APOE* variants. The substantial differences observed in comparison to *APOE-KO* indicate a gain-of-function mechanism by *APOE4* rather than a loss-of-function.

## 2. Materials and Methods

### iPS cell culture

*APOE*-isogenic iPS cell lines BIONi10-C3 (*APOE-KO*), BIONi10-C6 (*APOE2*), BIONi10-C2 (*APOE3*), and BIONi10-C4 (*APOE4*) (Schmid et al., 2019, 2020; de Leeuw et al., 2022) have been purchased from EBiSC. iPSCs were cultured on vitronectin (1:25) (100-0763, StemCell Technologies) coated plates in mTESR+ medium (100-0276, StemCell Technologies), split every 3-4 days and culture medium was exchanged every other day. For splitting, cells were washed with DPBS, incubated with ReLeSR (5872, StemCell Technologies) for 4 min at 37°C with 5% CO2, detached in 1mL mTESR+ and transferred to a new plate.

### iAstrocyte differentiation

iAstrocyte differentiation was carried out as described earlier (de Leeuw et al., 2022). In brief, iPSC lines were differentiated into NPCs by applying dual SMAD inhibition. On day 0, cells (appr. 20% confluent) were washed with PBS and medium changed to Neural Induction Medium 1, including 50% advanced DMEM/F12 (21331020, Gibco), 50% Neurobasal (21103-049, Gibco), 1x N2 (17502048, Thermo scientific), 1x B27 (17504044, Thermo Scientific), 2 mM Glutamax (Gibco) and 10 ng/mL hLIF (AF-300-05, Peprotech), 4 µM CHIR99021 (130-103-926, Miltenyi), 3 µM SB431542 (130-106-543, Miltenyi), 2 µM Dorsomorphin (130-104-466, Miltenyi), and 0.1 µM Compound E (73952, StemCell Technologies). Medium was changed daily. On day 3, cells were washed with PBS and medium was changed to Neural Induction Medium 2, including 50% advanced DMEM/F12, 50% Neurobasal, 1x N2, 1x B27, 2 mM Glutamax, 10 ng/mL hLIF, 4 mM CHIR99021, 3 mM SB431542, and 0.1 mM Compound E. On day 7, cells (appr. 80% confluent) were passaged from 12 WP to 6WP coated with 15 mg/mL poly-L-ornithine (P4957, Sigma-Aldrich) and 10 mg/mL laminin (L2020, Sigma-Aldrich). The medium was changed to neural stem cell maintenance medium (NSMM), including 50% advanced DMEM/ F12, 50% Neurobasal, 1x N2, 1x B27, 2 mM Glutamax, 10 ng/mL hLIF, 3 µM CHIR99021, and 2 µM SB431542. Medium was changed every day and cells were passaged twice a week (at 90-100% confluency). The first five passages 2 µM Thiazovivin (SML1045, Sigma) was added and after five passages 5 ng/mL FGF (PHG6015, Thermo Fisher) and 5 ng/mL epidermal growth factor (EGF) (PHG6045, Thermo Fisher) was added.

NPC passages 1-4 were further differentiated to astrocytes replating them onto 1 mg/mL Fibronectin (F1141-1MG, Sigma-Aldrich) (135.000 cells/well in 6WP) coated plates and changing the medium one day later to complete astrocyte medium (AM) (1801, ScienCell) supplemented with 2% FCS (0010, ScienCell) and AGS (1852, ScienCell) and P/S (0503, ScienCell) for 30 days. Medium was changed every other day and cells were passaged at 80%– 100% confluency. For the main stock, cells were frozen in BAMBANKER serum-free cryopreservation medium (WAKO302-14681, Avantor) at d28 which was the last time point before maturation of the astrocytes. For maturation and elimination of remaining proliferative cells, astrocytes were replated (350.000 cells/well in 6WP) and medium was changed to AM with 2 mM AraC (C1768, Sigma) but without FCS and then changed every other day. From day 37, cells were cultured in AM without FCS and without AraC until day 45. Cells were always replated at least 24h before an experiment.

### Immunocytochemistry

iAstrocytes were fixated after differentiation at d45 for 20 minutes at room temperature (RT) with 4% Paraformaldehyde (PFA) (47377.9L, VWR) and 4% sucrose (S9378, Merck) in DPBS. After washing the cells three times with PBS for about 5 minutes at RT, cells were blocked with 10% donkey serum (D9663, Sigma Aldrich) and 0.1% Triton (X100, Sigma Aldrich) in DPBS for 1h at RT. After washing with DPBS, cells were incubated with primary antibodies diluted in 3% donkey serum and 0.1% Triton in DPBS overnight at 4°C. After washing at the next day, secondary antibodies diluted in 3% donkey serum and 0.1% Triton in DPBS were added to the cells and incubated for 2h at RT in the dark. After washing, cells were incubated with 0.4 ng/µl DAPI (D9542, Sigma Aldrich) diluted in DPBS for 10 minutes. After another washing step, the coverslips were mounted with Mowiol (81381, Sigma Aldrich) on microscope objectives and stored over night at 4°C protected from light.

For calculation of differentiation efficiency, the number of s100β-positive cells was calculated per total DAPI count.

| *Primary antibody* | *Producer* | *Cat. No.* | *Dilution* |
| --- | --- | --- | --- |
| Anti-S100β | Sigma | S2532 | 1:200 |
| Anti-GJA1 | Abcam | Ab235585 | 1:200 |

| *Secondary antibody* | *Producer* | *Cat. No.* | *Dilution* |
| --- | --- | --- | --- |
| Dk-α-ms-cy3 | Jackson ImmunoResearch | 715-165-151 | 1:1000 |
| Dk-α-rb-cy5 | Jackson ImmunoResearch | 715-175-152 | 1:1000 |

### Protein extraction

iAstrocytes were harvested with Accutase. Cells were resuspended in RIPA buffer supplemented with protease inhibitor (11697498001, Sigma Aldrich). Three cycles of 30s of sonication were used to disrupt the cellular membranes. To extract the proteins, samples were centrifuged at 20.000g for 10 min at 4°C and the supernatant was finally collected and stored at −20°C until usage. Protein concentrations were determined with the Pierce BCA Assay Kit (23252, Thermo Fisher Scientific) according to the manufacturer’s instructions and the absorption at 562 nm was measured with the Infinite M Nano plate reader (Tecan).

### Immunoblotting

Each sample was diluted in RIPA buffer to obtain a concentration of 10 µg and mixed with sample buffer (NP0007, Thermo Fisher Scientific). Samples were denatured for 5 min at 95°C. Seeblue2 plus protein ladder (LC5925, Thermo Fisher Scientific) and samples were loaded onto 10-20% Tricine SDS-PAGE gels (EC6625BOX, Invitrogen) and run at 60 V for 15 min and 100 V for 90 min. Blotting was performed with the Trans-Blot Turbo Mini 0.2 µm nitrocellulose Transfer Pack (1704158, Bio-Rad) and the Trans-Blot Turbo Transfer System (1704158, Bio-Rad) at 2.5 A with 25 V for 7 min. Membranes were washed with 0.05% Tween (P1379, Sigma Aldrich) in PBS and blocked with 5% milk solution (A0830, ITW Reagents) in PBS for 1h at RT shaking. Membranes were then washed three times with PBS-Tween and incubated with primary antibodies diluted in 5% milk solution in PBS-Tween overnight at 4°C shaking. The next day, membranes were washed three times with PBS-Tween and incubated with secondary antibodies diluted in 5% milk solution in PBS-Tween for 2h at RT in the dark shaking. Membranes were washed three times, developed with one of the ECL selection kits (RPN2232/ RPN2235, Cytiva; 32106, Thermo Fisher Scientific) and imaged at the Image Quant 800 (Cytiva). Background subtraction was performed, and protein expressions were normalized to the housekeeping marker GAPDH/β-actin.

| *Primary antibody* | *Producer* | *Cat. No.* | *Dilution* |
| --- | --- | --- | --- |
| Anti-MFN1 | Cell Signaling | 14739S | 1:250 |
| Anti-MFN2 | Cell Signaling | 11925S | 1:1000 |
| Anti-FIS1 | Abcam | ab15686 | 1:10.000 |
| Anti-OPA1 | Cell Signaling | 67589S | 1:2000 |
| Anti-GAPDH | Meridian | H86504M | 1:5000 |
| Anti-hexokinase 1 | Abcam | ab154839 | 1:1000 |
| Anti-hexokinase 2 | Abcam | ab191838 | 1:1000 |

| *Secondary antibody* | *Producer* | *Cat. No.* | *Dilution* |
| --- | --- | --- | --- |
| Dk-α-gt-peroxidase | Jackson ImmunoResearch | 705-036-147 | 1:5000 |
| Dk-α-ms-peroxidase | Jackson ImmunoResearch | 715-035-151 | 1:5000 |
| Dk-α-rb-peroxidase | Jackson ImmunoResearch | 111-035-144 | 1:5000 |

### Seahorse assay

Using the Seahorse XFe24 (Agilent) the oxygen consumption rate (OCR) and the extracellular acidification rate (ECAR) were measured. Based on these measurements, all other factors were calculated. Seahorse experiments were performed according to the manufacturer’s instructions. Sensors were preincubated in H2O at 37°C without CO_2_ overnight and changed to calibrant 1h before the assay. Optimal cell density for each cell type was tested in prior Seahorse experiments. iAstrocytes were replated at 40.000 cells/well the day before the assay which was performed at day 45 or day 46 of differentiation. For each assay four measurements at baseline and three measurements after drug induction were performed. Each assay type was repeated three times for each cell line and included four background wells for each assay. The Seahorse DMEM medium (103575-100, Agilent) was freshly supplemented with 1 mM pyruvate and 2 mM L-glutamine and for the ATP rate and Mito Stress Test also with 10 mM glucose.

For the ATP Rate Assay drugs were used at the final concentrations of 1.5 µM oligomycin and 0.5 µM Rotenone + Antimycin A (ROT/AA). For the Mito Stress Test drugs were used at the final concentrations of 1.5 µM oligomycin, 2 µM Carbonyl cyanide-4 phenylhydrazone (FCCP) for and 0.5 µM Rotenone + Antimycin A (ROT/AA). FCCP titration was done in earlier Seahorse experiments. For the Glyco Stress Test drugs were used at a final concentration of 10 mM glucose, 1 µM oligomycin and 50 mM 2-deoxy-glucose (2-DG).

After Seahorse assays, cells were immediately fixated with 4% PFA and 4% sucrose in DPBS and stained with 0.4 ng/µL DAPI for 10 min at RT. DAPI was imaged with an inverted fluorescence microscope (Zeiss) with 10x magnification. The number of nuclei in each well was automatically analyzed by a macro written in Fiji and used for normalization of the Seahorse data with the Wave Software.

### Fluorescence lifetime imaging (FLIM)

7-8 days before imaging, d37-39 iAstrocytes were replated onto microscope dishes (IBL, 220.110.012) at a density of 120.000 cells/dish. Cells were transduced with 1 µL AAV nanosensor 2h after replating, once the cells had begun to attach to the plate. The lactate sensor LiLac (Koveal et al., 2022) (ssAAV-2/2-shortCAG-6xHis_LiLac-WPRE-SV40p(A) was purchased from the viral vector facility of Neuroscience Center Zurich. After 24h, virus was removed with a medium change. Cells were normally cultured until the imaging experiment at day 45/46. Before imaging, cells were cultured for at least 1h in reference buffer. During imaging, microscope dishes were constant perfused with reference buffer containing 112 mM NaCl, 3 mM KCl, 1.25 mM CaCl_2_, 24 mM NaHCO_3_, 1.25 mM MgSO_4_, 10 mM HEPES, 2 mM glucose, 1 mM lactate, 0.1 mM pyruvate, 2 mM L-glutamine, at a rate of about 2 mL/min at a constant temperature around 34°C using a customized version of the PiFlow pump (Kassis et al., 2018). For a stable baseline, cells were given about 5-10 min to adjust to the setup. Drugs were added in addition to the reference buffer at a concentration of 5 mM sodium azide (Merck, 1.06688.0100) and 2 µM AR-C155858 (MedChem, HY-13248).

FLIM imaging was performed on a custom two-photon microscope (Mayrhofer et al., 2015) coupled to an Insight Deepsee (SpectraPhysics) femtosecond pulsed laser. A 25x water immersion objective (XLPlan N 25×/1.05w MP, WD = 2mm, Olympus) was used for image acquisition. The excitation and emission beam paths were separated using a dichroic mirror. Emission light was further divided into specific wavelength components with dichroic mirrors at 560 and 506 nm, then focused on a PMA Hybrid 40-mod HPD detector (Picoquant) equipped with filters for CFP (475/50;) and laser rejection (770SP). Data acquisition and system control were performed using a customized version of ScanImage 3.8 and custom LabVIEW software (Version 2012; National Instruments), while Time-Correlated Single-Photon Counting and FLIM image generation were performed using in-house instrumentation (Velasquez Moros et al., 2024). LiLac-expressing cells were excited at 870 nm and images were acquired at a 2.96 Hz with a 128×128 pixel resolution.

FLIM data analysis was conducted using FLIManalysis (https://gitlab.com/einlabzurich/flimanalysis), a Matlab-based wrapper of the FLIMfit library (Warren et al., 2013) designed to facilitate batch processing. ROIs were selected using ImageJ and including ROI-based decay summation was performed before fitting.

### Proteomics

For this study, we analyzed a proteomic dataset that we previously generated (de Leeuw et al., 2022) and focused on yet unpublished pathways related to energy metabolism. The detailed methodology for sample preparation, mass spectrometry acquisition and data processing has been described (de Leeuw et al., 2022). Briefly, iAstrocytes were cultured, treated and lysed in RIPA buffer before protein extraction using the iST Kit (PreOmics, Germany). Mass spectrometry analysis was performed using an Orbitrap Fusion Lumos (Thermo Scientific) coupled to an M-Class UPLC (Waters).

For the present study, we focused on proteins and pathways associated with cellular energy metabolism. The acquired raw MS data were reprocessed using MaxQuant and statistical analyses were performed in R. We conducted pathway enrichment analysis using the fgsea R/Bioconductor package, leveraging gene sets from the Molecular Signatures Database (MSigDB). Protein lists were ranked based on moderated t-statistics and enrichment scores were computed to identify significantly altered pathways. All data processing steps were conducted as previously described, with modifications specific to the analysis of metabolic pathways.

### Metabolomics - Sample preparation for LC-MS analysis of polar metabolites

iAstrocytes were cultured until day 45. Following a single PBS wash, polar metabolites were extracted using a MeOH:ACN:H2O (40:40:20) solution. The plate was manually shaken for 30 seconds and incubated at −80°C with closed lids for 15 minutes. Cells were then detached using a cell scraper, transferred to Eppendorf tubes and stored at −80°C. For each cell line three batches were collected before samples being delivered to the Functional Genomics Center Zurich. For each sample, 4 ml of extract was delivered to the Functional Genomic Center Zürich to perform LC-MS based untargeted analysis of the polar metabolites content. The extracts have been centrifuged for 20 min at 7000 rpm and 4°C to precipitate proteins and cell debris, 3 ml of clear supernatant was progressively transferred to clean 2 ml test tubes and dried under nitrogen flow prior to solubilization of the metabolites in injection solvent (90% acetonitrile). The protein pellet was used to determine the protein content of each sample, and the protein content was later used for data normalization.

Before measuring, the samples were centrifuged for 10 minutes at 10.000 rpm/ 4°C and the clear supernatant was transferred to glass vials suitable for LC-MS analysis (Total Recovery Vials, Waters). In addition, method blanks, mixtures of pure standards, and pooled samples were prepared in the same way to serve as quality control for the measurements.

### Metabolomics - LC-MS data acquisition

Metabolites were separated on a Thermo Vanquish Horizon Binary Pump equipped with Waters Premier BEH Amide column (150 mm x 2.1 mm), applying a gradient of 10 mM ammonium bicarbonate in 5% acetonitrile pH9 (A) and 10 mM ammonium bicarbonate in 95% acetonitrile (B) from 99% B to 30% B over 12 min. The injection volume was 5 μl while the flow rate was 0.4 ul/min with column temperature of 40°C and autosampler temperature of 5°C. The LC was coupled to Thermo Exploris 480 mass spectrometer by a HESI source. MS1 (molecular ion) and MS2 (fragment) data were acquired using negative polarization and Full MS / dd-MS² (Top5) over a mass range of 70 to 1050 m/z at MS1 and MS2 resolution of >17’500. Quality controls were run on pooled samples, reference compound mixtures, and blanks to determine technical accuracy and stability.

### Metabolomics - Untargeted Metabolomics Data analysis

The metabolomics dataset was evaluated in an untargeted fashion with Compound Discoverer software (Thermo Scientific). The modular data analysis workflow includes spectra selection, retention times alignment, compound detection and grouping, gap filling, background filtering and normalization (data are protein content normalized). mzCloud and mzVault have been used to score fragmentation patterns and assign MS2-based identities to the features. A filtering process was performed, leading to the manually annotated compound table, where each feature is annotated with the highest level of confidence. Filtering parameters used were the following: Signa/noise > 3, mzCloud or mzVault match >50, ppm mass error within +/− 5 ppm., match with in-house developed MS1_RT library within +/− 10 sec, chromatographic peak and MS2 spectra quality.

Enrichment analysis was performed using Metaboanalyst 6.0 (Pang et al., 2024). For comparison of two groups the quantitative enrichment analysis function was selected, pathways were identified according to small molecule pathway database (SMDB) library and results were plotted using R.

### Statistics

Normal distribution was tested by using the Shapiro-Wilk normality test and the Kolmogorov-Smirnov test. Outliers were identified with the ROUT test (Q = 1%). Differences between groups were analyzed by one-way ANOVA followed by Tukey test for multiple comparisons for normally distributed data or by the Kruskal-Wallis test followed by Dunn’s multiple comparisons test for not normally distributed data. For the Seahorse assays, 3 - 5 wells per cell line with each containing about 40.000 cells were analyzed per assay and each assay was repeated at least 3 times independently. For FLIM 1 dish containing 120.000 cells of which 10 were analyzed per experiment, which were repeated at least 4 times independently for each cell line. For MS-based proteomics and metabolomics, 3 biological replicates were analyzed per genotype.

## 3. Results

### 3.1. APOE4 decreases ATP production based on glycolysis and mitochondrial respiration

To determine the *APOE* effect on human astrocytes, we differentiated APOE-isogenic iPS cell lines (*APOE-KO*, *APOE2*, *APOE3* and *APOE4*) into functional iAstrocytes **(Fig. 1A)**. Both the *APOE*-isogenic iPSCs and iAstrocytes had already been characterized and functionally validated in previous studies (de Leeuw et al., 2022; Budny et al., 2024). To verify successful iAstrocytes differentiation, immunostaining was performed confirming the expression of astrocyte markers s100β and GJA1 **(Fig. S1A)**. Differentiation efficiency was calculated as s100β-positive cells per total DAPI counts and reached >97% for all cell lines (**Fig. S1B**). To determine whether *APOE* genotypes differentially regulate the glycolytic and mitochondrial ATP production in our *APOE*-isogenic iAstrocytes, a Seahorse ATP rate assay was performed. Oxygen consumption rate (OCR) and extracellular acidification rate (ECAR) were measured four times at baseline and three times after each drug administration **(Fig. 1B, C)**. Oligomycin and Rotenone/Antimycin A (ROT/AA) were used to inhibit respiratory complex V and I/III, respectively, based on which mitochondrial and glycolytic ATP production could be analyzed. Mitochondrial ATP production was significantly lower in *APOE4* iAstrocytes than in all other *APOE* lines (E4<E3=E2=KO) **(Fig. 1D)**. ATP production based on glycolysis as well as total ATP production was lower in *APOE*3 and *APOE4* iAstrocytes compared to *APOE2* and *APOE-KO* (E4=E3<E2=KO) **(Fig. 1E, F)**. The lower total ATP production in *APOE3* compared to *APOE2* and *APOE-KO* iAstrocytes was mainly caused by a reduction in glycolysis rather than in mitochondrial respiration. Interestingly, the proportion of glycolsis-generated ATP was higher *APOE2* and *APOE-KO* iAstrocytes (41% and 42%, respectively) than in *APOE3* (29%) and *APOE4* iAstrocytes (33%). **(Fig. 1G)**.

**Fig. 1:**
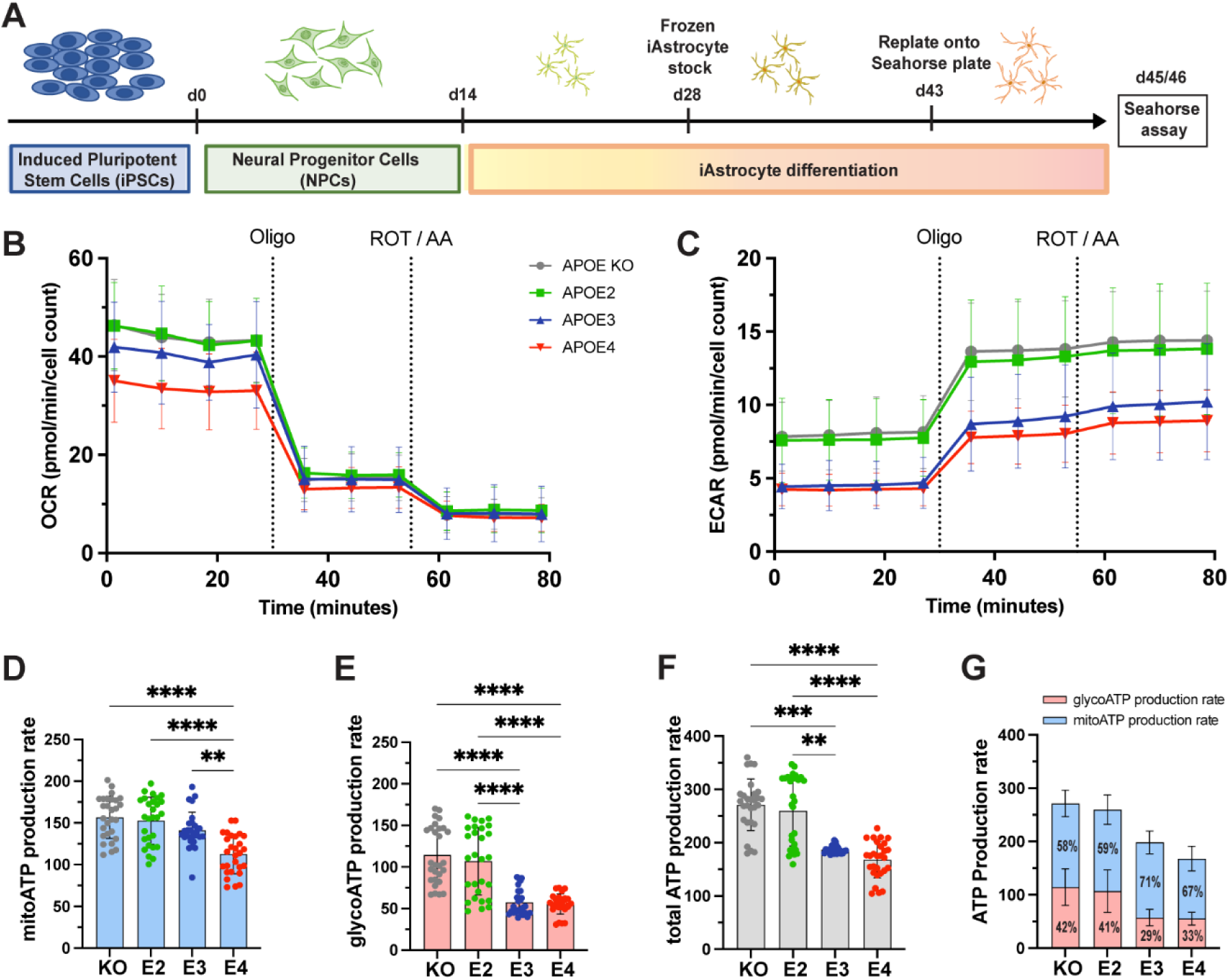
ATP production in *APOE*-isogenic astrocytes. **A:** Schematic differentiation timeline of *APOE*-isogenic iAstrocytes for the Seahorse assays. **B:** OCR and **C:** ECAR were measured in iAstrocytes. *APOE-KO* in grey, *APOE2* in green, *APOE3* in blue and *APOE4* in red. Oligo: Oligomycin; ROT/AA: Rotenone/Antimycin A. **D:** Mitochondrial ATP (mitoATP) production rate, **E:** glycolytic ATP (glycoATP) production rate and **F, G:** total ATP production rate, calculated based on OCR and ECAR measurements. glycoATP in red and mitoATP in blue. Data was analyzed using the Kruskal-Wallis test (Dunn’s multiple comparisons test) (D-F). ATP rate assay was repeated seven times independently. ** p < 0.01, *** p < 0.001, **** p < 0.0001.

### 3.2. APOE4 affects mitochondrial respiration and increases mitochondrial stress

To gain a deeper understanding of how the different *APOE* genotypes affect oxidative energy metabolism, we made use of data previously generated by an unlabeled mass spectrometry-based proteomic screen (de Leeuw et al., 2022). Gene set enrichment analysis (GSEA) of yet unpublished proteomic data showed strong upregulation of pathways involved in OXPHOS as well as mitochondrial structure and function in *APOE4* compared to *APOE2* and a tendency to downregulation in *APOE2* compared to *APOE3* **(Fig. 2A)**. It should be noted that the *APOE-KO* line had not been included in the proteomic analysis, thus proteomic data from this line was not available.

**Fig. 2:**
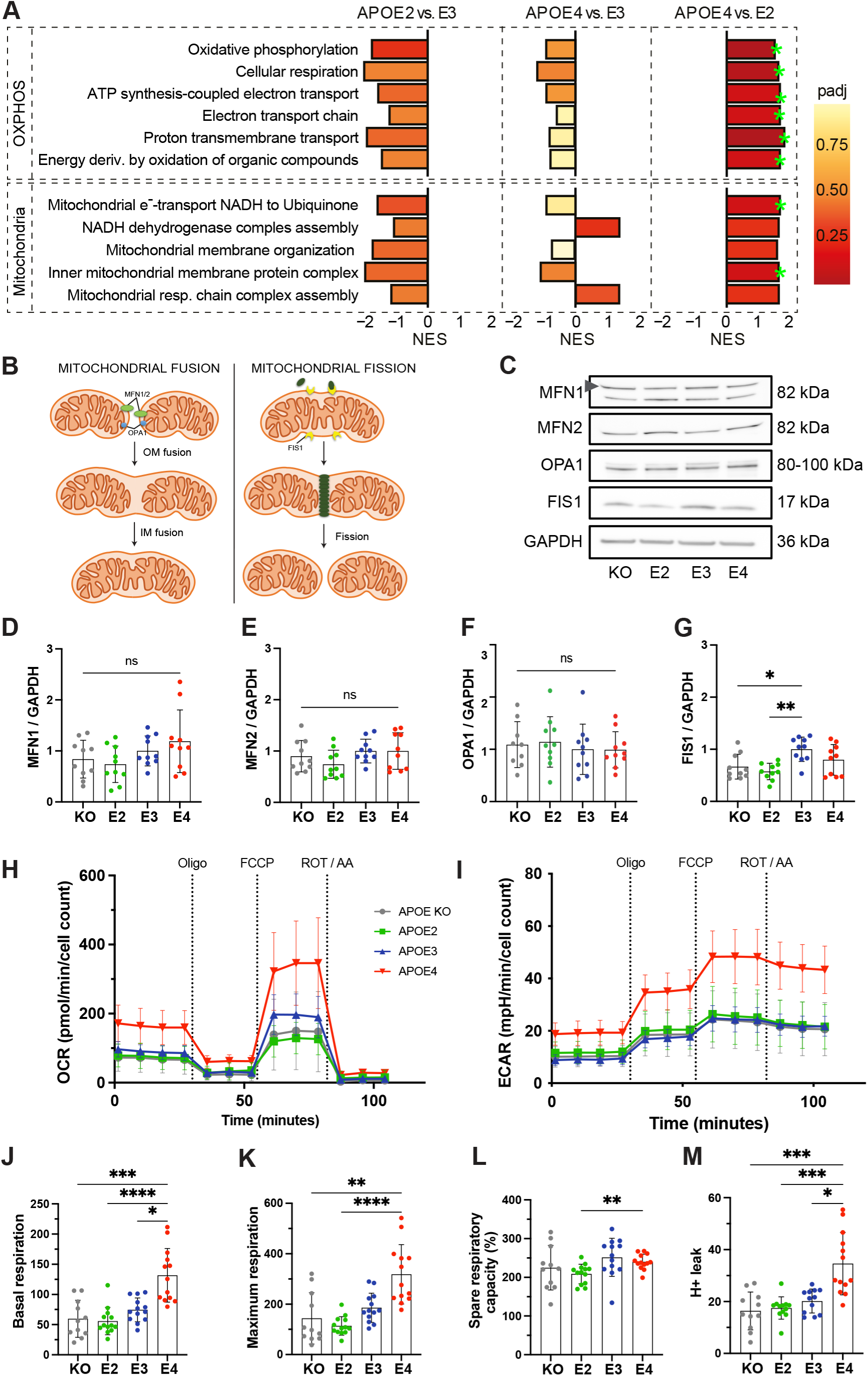
Mitochondrial function in *APOE*-isogenic iAstrocytes. **A:** GSEA normalized enrichment scores (NESs) of proteomic lists ranked according to the t-statistic obtained for the contrast *APOE2* versus *APOE3*, *APOE4* versus *APOE3*, and *APOE4* versus *APOE2* iAstrocytes. NES is plotted on the x axis, with color-coded bars for the individual gene ontology (GO) terms. Gene sets with adjusted p-values <0.2 are annotated (green stars). **B:** Schematic overview of mitochondrial fusion and fission markers. **C:** Representative western blot images of MFN1, MFN2, FIS1 and OPA1 in APOE-KO, -E2, -E3 and-E4 isogenic iAstrocytes. **D-G:** Quantified protein levels of MFN1, MFN2, FIS1 and OPA1, normalized to GAPDH. **H:** OCR and **I:** ECAR measurements. *APOE-KO* in grey, *APOE2* in green, *APOE3* in blue and *APOE4* in red. Oligo: Oligomycin; FCCP: Carbonyl cyanide-4 (trifluoromethoxy) phenylhydrazone; ROT/AA: Rotenone/Antimycin A. **J:** Basal respiration, **K:** maximal respiration, **L:** spare respiratory capacity and **M:** proton leak in iAstrocytes. Data was analyzed using the Kruskal-Wallis test (Dunn’s multiple comparisons test) (J, K, M) or one-way ANOVA (Tukey test for multiple comparisons) (D-G, L). Western blots were repeated ten times and Mito Stress Test was repeated three times independently. ns = not significant, * p < 0.05, ** p < 0.01, *** p < 0.001, **** p < 0.0001.

Efficient energy production relies on maintaining a healthy and functional mitochondrial network, which is regulated by dynamic reshaping events known as fusion and fission **(Fig. 2B)**. To determine whether these processes are affected by the APOE genotype, proteins associated with mitochondrial fusion and fission were analyzed **(Fig. 2C-G)**. No differences in the mitochondrial fusion proteins mitofusin-1/2 (MFN1/2) and OPA1 were detected between the *APOE*-isogenic lines **(Fig. 2D-F)**. However, levels of mitochondrial fission marker FIS1 were higher in *APOE3* compared to *APOE-KO* and *APOE2* **(Fig. 2G)**.

While ATP production based on mitochondrial respiration was reduced in *APOE4* **(Fig. 1D)**, proteomic data indicated an increase in pathways related to oxidative phosphorylation **(Fig. 2A)**. To further dissect the *APOE4* effect on mitochondrial function, basal and maximal respiration as well as mitochondrial capacity and proton leak were analyzed using the Seahorse Mito Stress Test. OCR and ECAR were measured both at baseline and after the addition of Oligomycin, carbonyl cyanide-4 (trifluoromethoxy) phenylhydrazone (FCCP) and ROT/AA **(Fig. 2H, I)**. FCCP was used to induce stress, which collapses the proton gradient and disrupts the mitochondrial membrane potential. This disruption allows for unrestricted electron flow through the electron transport chain, ultimately leading to maximum oxygen consumption at complex IV. The OCR values were then used to calculate mitochondrial properties. Basal and maximal respiration was significantly higher in *APOE4* iAstrocytes compared to the genotypes **(Fig. 2J, K)**, while spare respiration capacity was increased in *APOE4* compared to *APOE2* **(Fig. 2L)**. Further, a significantly elevated proton leak was observed in *APOE4* iAstrocytes compared to all other lines **(Fig. 2M).** This indicates that *APOE4* increases mitochondrial respiration and causes mitochondrial dysfunction.

### 3.3. APOE4 increases glycolysis

Since *APOE4* did not only affect mitochondrial ATP production but also decreased ATP production based on glycolysis compared to *APOE-KO* and *APOE2*, we further assessed glycolytic function in iAstrocytes. Protein levels of the first enzymes in the glycolytic pathway, hexokinase 1 and 2 (HK1, HK2), were measured **(Fig. 3A)**. No differences in HK1 were observed (**Fig. 3B**), while protein levels of HK2 were significantly lower in *APOE3* and *APOE4* compared to *APOE-KO* and *APOE2* (KO=E2>E3=E4) **(Fig. 3C)**. This is in agreement with the reduction of glycoATP production in *APOE3* and *APOE4* (**Fig. 1E**). Subsequently, the Seahorse Glyco Stress Test was performed to tease out effects glycolytic function, glycolysis as well as glycolytic capacity and reserve. OCR and ECAR were measured both at baseline and after the addition of glucose, Oligomycin and 2-deoxy-glucose (2-DG) **(Fig. 3D, E)**. 2-DG is a glucose analog that inhibits glycolysis by competitively binding HK. Based on ECAR values, glycolytic properties were analyzed. *APOE4* astrocytes exhibited increased glycolysis and glycolytic capacity compared to *APOE-KO* and *APOE2* **(Fig. 3F, G)**, while glycolytic reserve remained unchanged between *APOE* lines **(Fig. 3H)**. These results indicate that *APOE4* affects glycolytic function on both protein and functional level.

**Fig. 3:**
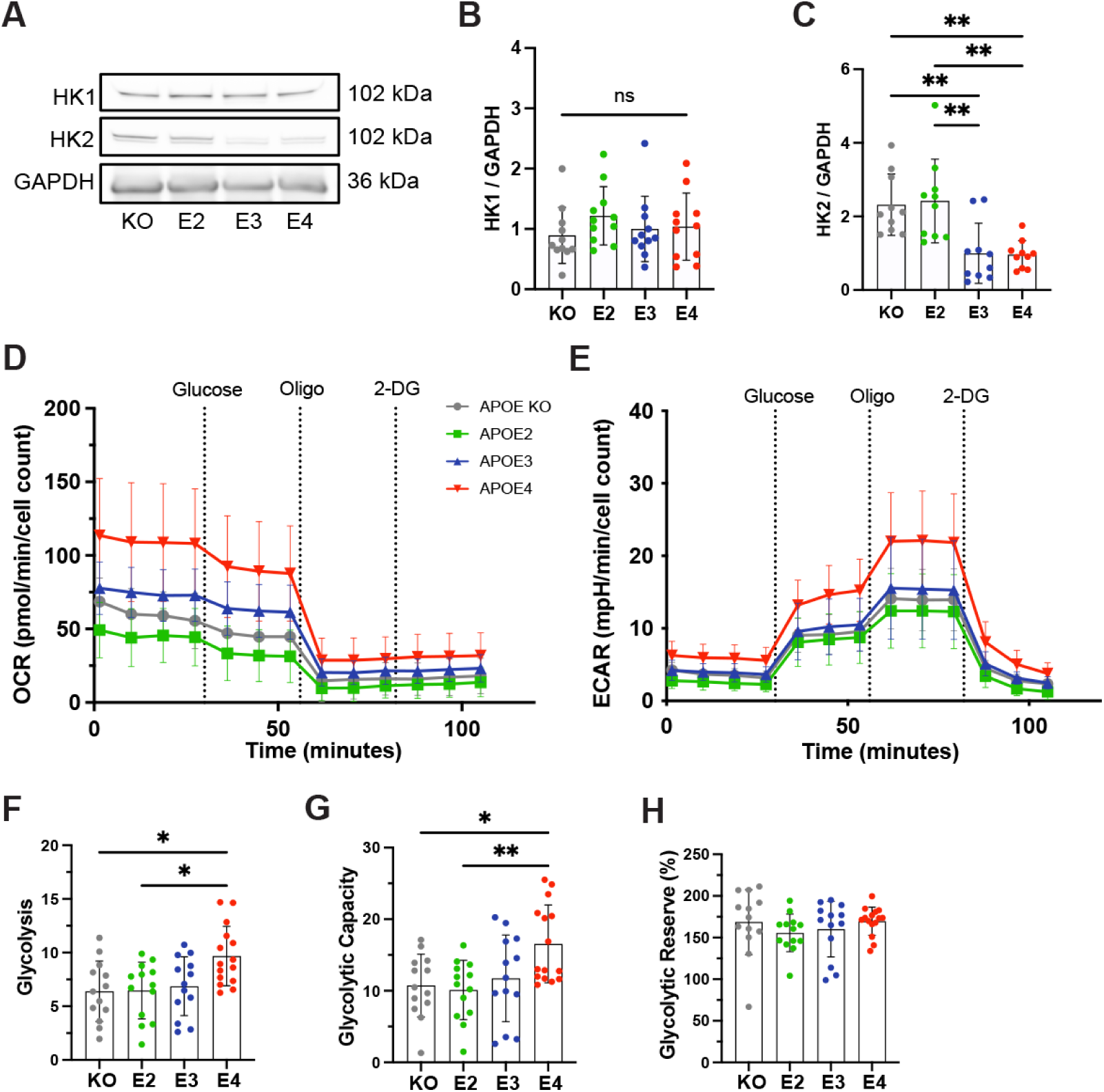
Glycolytic function in *APOE*-isogenic iAstrocytes. **A:** Representative western blots of HK1, HK2 and GAPDH in *APOE-KO*, -*E2*, -*E3* and-*E4* iAstrocytes. **B, C:** Quantified protein levels of HK1 and HK2 in *APOE-KO*, -*E2*, *E3* and-*E4* iAstrocytes. All protein levels were normalized to GAPDH. **D:** OCR and **E:** ECAR in iAstrocytes. *APOE-KO* in grey, *APOE2* in green, *APOE3* in blue and *APOE4* in red. Oligo: Oligomycin; Glu: Glucose; 2-DG: Glucose analog 2-deoxy-D-glucose. **F:** Glycolytic function, **G:** glycolytic capacity and **H:** glycolytic reserve in iAstrocytes. Data was analyzed using the Kruskal-Wallis test (Dunn’s multiple comparisons test) (C, H) or one-way ANOVA (Tukey test for multiple comparisons) (B, F, G). Western blots were repeated at least ten times and Glyco Stress Test was repeated three times independently. ns = not significant, * p < 0.05, ** p < 0.01.

### 3.4. APOE genotype does not affect lactate dynamics

Given the observed changes in glycolysis, we further investigated glycolytic function by analyzing lactate dynamics in more detail. iAstrocytes were transduced with LiLac, a nanosensor for lactate, and fluorescence lifetime microscopy (FLIM) imaging was used to analyze intracellular lactate levels (Koveal et al., 2022) **(Fig. 4A-C)**. Imaging was performed under cell culture-like conditions at 34°C and with carboxygenated reference buffer (RB) flow **(Fig. 4B)**. Basal lactate levels were measured during the perfusion with RB at the beginning of each experiment. No differences in basal lactate levels were observed between the lines **(Fig. 4D)**. After a constant baseline, 5 mM sodium azide was added to inhibit mitochondrial function and stimulate glycolysis. The rate of lactate increase was then assessed by slope analysis **(Fig. 4E; S2A-E)**. Similar rates of lactate increase through azide treatment were observed between *APOE* genotypes **(Fig. 4F)**. After a constant baseline again with RB, 2 µM of the monocarboxylate transport inhibitor AR-C was added **(Fig. 4G; S2A-E)**. The immediate decrease in measured lifetime is equivalent to an increase in intracellular lactate and shows that iAstrocytes predominantly produce lactate. By using slope analysis, the rate of lactate increases, which represents the rate of intracellular lactate accumulation after the inhibition of lactate export through monocarboxylate transporters was determined **(Fig. 4H)**. Similar rates of lactate increase through AR-C were observed between *APOE* lines. The Warburg index, a measure of the balance between glycolytic and oxidative metabolism, was then calculated based on dividing the AR-C slope through the azide slope. This indicates whether cells rely on aerobic glycolysis rather than mitochondrial respiration for energy production. While *APOE4* and *APOE3* showed a slightly reduced Warburg index, the difference between the genotypes was not significant **(Fig. 4I)**. Taken together, while *APOE* genotypes differentially affect mitochondrial respiration and glycolysis, no effect on lactate dynamics could be observed.

**Fig. 4:**
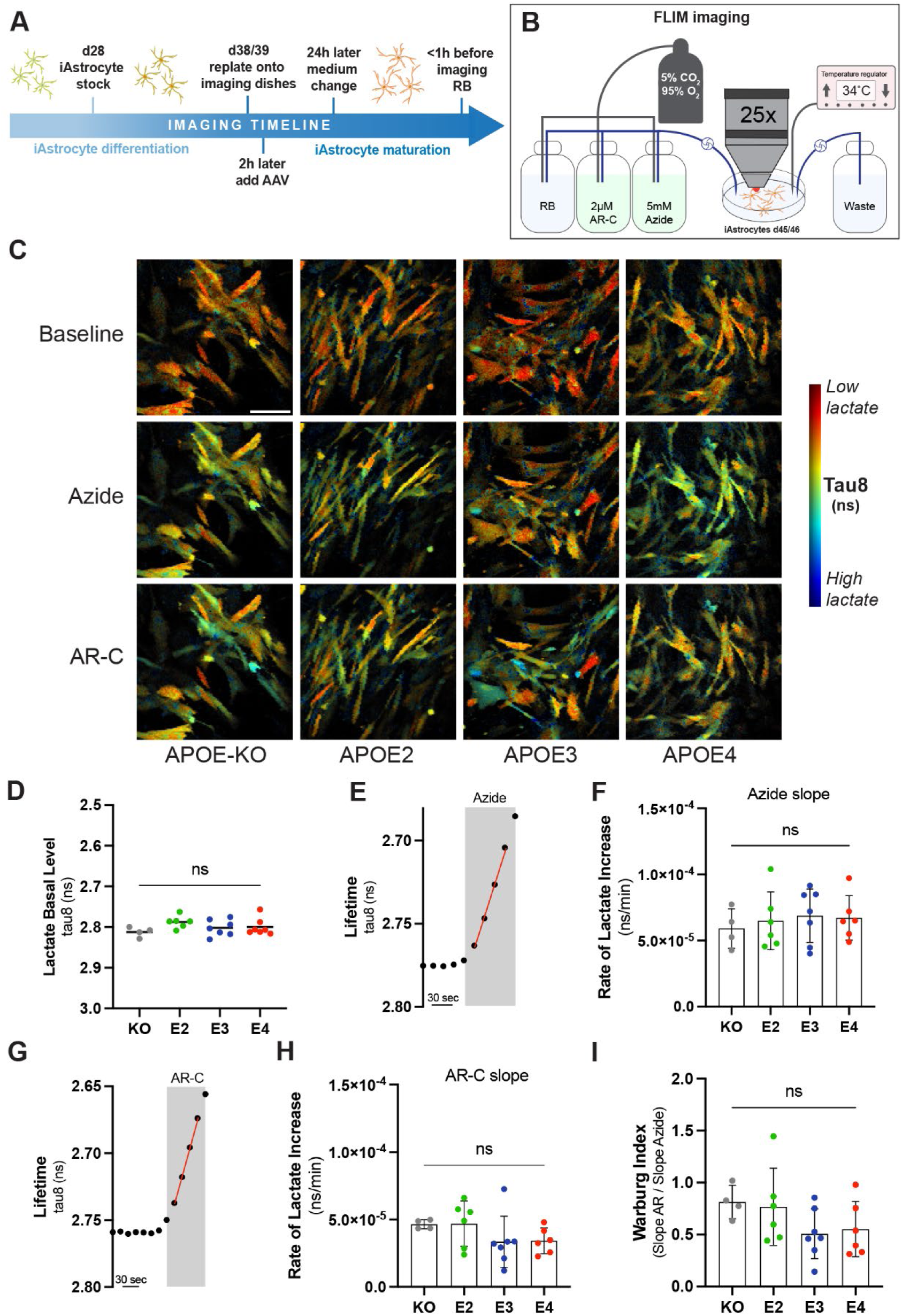
Lactate metabolism in *APOE*-isogenic iAstrocytes. **A:** Timeline of iAstrocytes preparation for FLIM imaging with the genetically-encoded lactate sensor LiLac. **B:** Schematic representation of FLIM setup. **C:** Representative intensity merged FLIM images in *APOE-KO*, -*E2*, -*E3* and-*E4* isogenic iAstrocytes at baseline and with Azide or AR-C treatment. In blue low tau8 (high lactate) and in red high tau8 (low lactate). Scale bar = 100 µM **D:** Quantification of tau8 (ns) for the basal lactate levels. **E:** FLIM protocol to measure the lifetime change in response to 5 mM Azide. An example measurement for *APOE3* is shown. **F:** Azide slop analysis for measuring the rate of intracellular lactate increase (ns/min). **G:** FLIM protocol to measure the lifetime change in response to 2 µM AR-C. An example measurement for *APOE3* is shown. **H:** AR-C slop analysis for measuring the rate of intracellular lactate increase (ns/min). **I:** Warburg analysis based on the Azide and AR-C slop analysis. Non-parametric data was analyzed using the Kruskal-Wallis test (multiple comparisons) (H) and parametric data was analyzed using the ordinary one-way ANOVA (multiple comparisons) (D, F, I). FLIM was repeated at least four times independently for each cell line. ns = not significant; RB = Reference buffer.

### 3.5. APOE genotype-dependent changes in energy metabolism-related pathways

Next, we applied liquid chromatography–mass spectrometry (LC-MS)-based metabolomic analysis of polar metabolites followed by enrichment analysis using MetaboAnalyst 6.0 to compute differentially regulated metabolic pathways in *APOE4* iAstrocytes compared to *APOE2*, *APOE3*, and *APOE-KO*. It should be noted that the enrichment ratio does not show whether a pathway is specifically enriched in one group over the other but rather indicates that a pathway is perturbed, with some metabolites being up- and others down-regulated due to dynamic regulation of metabolic flux. We specifically looked for pathways involved in cellular energy metabolism. Nine pathways related to glucose and energy metabolism were identified when comparing *APOE4* to *APOE2* iAstrocytes **(Fig. 5A),** while 7 pathways were identified in *APOE4* compared to *APOE3* **(Fig. 5B)** and 10 pathways in *APOE4* compared to *APOE-KO* **(Fig. 5C)**. Some pathways were only enriched in the comparison of *APOE4* to either *APOE2*, *APOE3* or *APOE-KO*, whereas others, such as the malate-aspartate-shuttle, glycolysis, mitochondrial electron transport chain, Warburg effect or pyruvate metabolism were found in all comparisons. However, the degree of pathway enrichment varied between the lines.

**Fig. 5:**
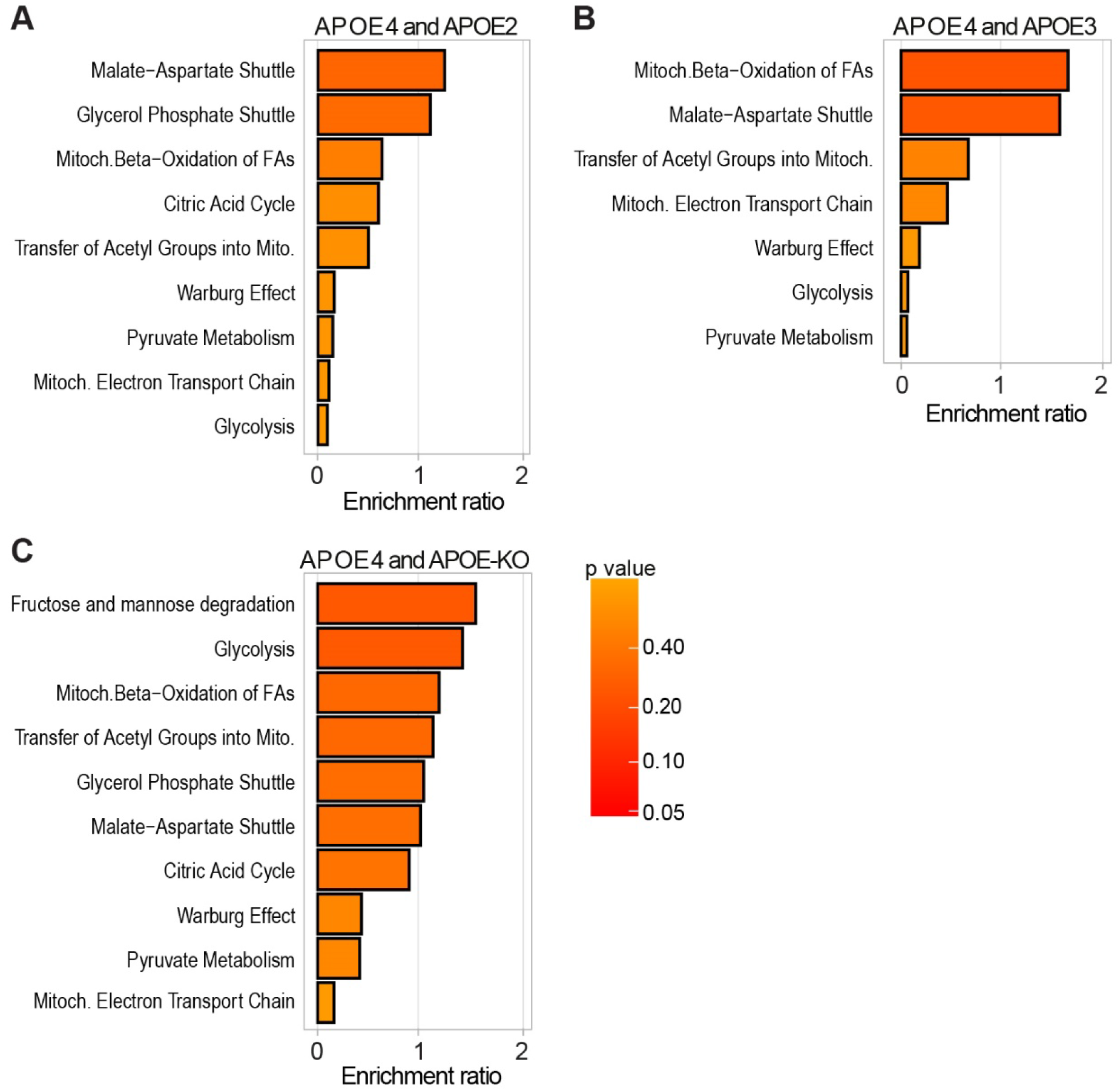
Metabolomic enrichment analysis in *APOE*-isogenic iAstrocytes. **A-C:** Enrichment analysis of glucose and energy metabolism-related pathways, compared between the respective *APOE*-isogenic iAstrocytes and analyzed using Metaboanalyst 6.0. Pathways are ranked according to the enrichment ratio.

When comparing *APOE4* to *APOE2*, the malate-aspartate shuttle ranked among the top 25 enriched metabolic pathways **(Fig. S3A)** and in the *APOE4* versus *APOE3* comparison mitochondrial beta-oxidation was among the top 25 enriched pathways **(Fig. S3B).** In *APOE4* compared to *APOE-KO*, fructose and mannose degradation and glycolysis were found in the top 25 pathways **(Fig. S3C)**.

In summary, LC-MS-based metabolomic analysis indicates that *APOE4* has a major impact on human astrocyte energy metabolism, confirming our observations from proteomic analysis and functional assays.

## 4. Discussion

AD has been increasingly linked to dysregulated energy metabolism, yet the underlying mechanisms remain unclear. Using *APOE*-isogenic iPSC-derived astrocytes, we investigated *APOE*-associated differences in metabolic function. Our model shows AD-relevant proteomic pathways being specifically upregulated in *APOE4* astrocytes and downregulated in *APOE2* (**Fig. S4**), highlighting the biological relevance of this system.

Our data demonstrates that *APOE4* expression leads to an increase in both mitochondrial respiration and glycolysis; however, this is accompanied by inefficient ATP production. These results align partially with existing literature, which however presents conflicting findings. While some studies reported reduced glycolysis in APOE4-expressing cells (Wu et al., 2018; Fang et al., 2021; Zhang et al., 2023), others indicated a metabolic shift from oxidative phosphorylation to glycolysis (Sonntag et al., 2017; Williams et al., 2020; Farmer et al., 2021; Lee et al., 2023), with *APOE4* astrocytes exhibiting greater glycolytic activity compared to *APOE3* astrocytes. Higher oxygen consumption together with an increased glycolysis while having reduced ATP production is a sign of mitochondrial uncoupling. This occurs when the electron transport chain (ETC) continues to consume oxygen and generate a proton gradient, but ATP synthesis is less efficient because protons leak back into the mitochondrial matrix without driving ATP synthase. It leads to increased mitochondrial activity, including higher oxygen consumption as well as to increased glycolysis to compensate for ATP loss, but lower ATP production (Cadenas, 2018; Demine et al., 2019).

A potential mitochondrial uncoupling caused by *APOE4* is supported by our finding that *APOE4* iAstrocytes show a significant proton leak. Changes in proton leak are an indicator for mitochondrial damage such as mitochondrial uncoupling. Increased mitochondrial respiration together with decreased ATP levels can also be induced by treating cells with mitochondrial uncouplers supporting our hypothesis of *APOE4* may induce mitochondrial uncoupling (Demine et al., 2019; Shrestha et al., 2021). In the present study, we did not observe any changes in mitochondrial fusion and fission proteins due to *APOE* genotype, which is consistent with our previous findings in AD patient-derived cells as well as *APOE*-isogenic neurons (Birnbaum et al., 2018; Budny et al., 2024). This suggests that while mitochondrial function is altered by *APOE*, these changes are not driven by modifications in mitochondrial fission or fusion. Recent studies have shown the role of mitochondrial uncoupling proteins (UCPs) in AD. UCP4 levels are significantly reduced in AD brain tissue and the UCP4 variant rs9472817 increases AD risk, especially in *APOE4* carriers, suggesting a connection between UCP4 and *APOE4* in disease progression (de la Monte and Wands, 2006; Montesanto et al., 2016). Overexpressing UCP4 in astrocytes in AD models prevents mitochondrial dysfunction and improves memory (Rosenberg et al., 2023). Mitochondrial dysfunction and oxidative stress, early features of AD, are linked to UCP4 downregulation due to inflammation (Thangavel et al., 2017). These studies support the role of mitochondrial dysfunction in AD pathology and particularly in relation to *APOE4*.

We show that *APOE4* iAstrocytes exhibit lower expression of HK2, which may be attributed to mitochondrial dysfunctions. HK2 is typically associated with the outer mitochondrial membrane (OMM) through its interaction with the outer mitochondrial voltage-dependent anion channel (VDAC) (Haloi et al., 2021). Disruption of this interaction, possibly due to changes in mitochondrial membrane potential or metabolic flux, could result in the dissociation of HK2 from the membrane, leading to its downregulation. Notably, VDAC has been considered a potential target for AD, as it plays a crucial role in mitochondrial function and VDAC malfunction can result in impaired energy production, increased oxidative stress and mitochondrial homeostasis disruption (Fang and Maldonado, 2018; Yang et al., 2024). Interestingly, HK1 expression remained unchanged in our model. This may suggest that HK1 is not as significantly impacted by mitochondrial changes in *APOE4* iAstrocytes and its expression might remain stable due to its involvement in maintaining basal glycolytic function and supporting alternative pathways like the pentose phosphate pathway (PPP), rather than rapid glycolysis for ATP production. Our findings are consistent with previous studies demonstrating significant differences in HK2 protein expression between *APOE4* and *APOE2*. However, other studies have also reported lower HK2 expression in *APOE4* compared to *APOE3*, a difference we did not observe (Wu et al., 2018; Zhang et al., 2023). This discrepancy may be attributed to the fact that HK2 expression differences between *APOE* variants become more pronounced with increasing passage numbers (Zhang et al., 2023). Similarly, Fang and colleagues found that the effects of *APOE* on OCR and ECAR in *APOE4* astrocytes were not prominent after one month but became evident after two months of culture (Fang et al., 2021). Together, these findings suggest a potential metabolic shift in *APOE4* astrocytes that intensifies over time, highlighting the importance of the time point of analysis when comparing results.

Further, *APOE-KO* and *APOE2* iAstrocytes showed higher glycoATP production and higher FIS1 and HK2 protein expression not only compared to *APOE4* but also to *APOE3*. A noticeable trend is observed in the glyco/mitoATP production ratio, with glycolytic ATP production being higher in *APOE-KO* and *APOE2* compared to *APOE3* and *APOE4*. This indicates that *APOE-KO* and *APOE2* cells rely more on glycolysis compared to *APOE3*. The study by Wu and colleagues showed similar findings, as differentiated N2a cells expressing APOE2 exhibited the highest HK expression and glycolytic activity. This suggests that *APOE-KO* and *APOE2* may promote a more robust glycolytic energy profile compared to *APOE3* cells (Wu et al., 2018; Zhang et al., 2023).

This study represents the first application of a genetically encoded nanosensor for lactate in *APOE*-isogenic cells, providing novel insights into intracellular lactate metabolism. Our findings reveal similar basal levels of intracellular lactate across all *APOE* variants, with no significant differences in lactate accumulation rates or Warburg index between the *APOE* variants. Notably, our iAstrocytes exhibit a Warburg index comparable to that previously reported (San Martín et al., 2013). These results suggest that mitochondrial dysfunction, as evidenced by the proton leak, may enhance oxygen consumption and overall energy expenditure. However, rather than leading to excessive lactate accumulation, glycolysis appears to be upregulated just enough to maintain metabolic balance. This implies a compensatory mechanism where pyruvate is preferentially directed toward mitochondrial oxidation rather than lactate production, thereby preserving stable intracellular lactate dynamics. Interestingly, *APOE3* cells showed reduced glycoATP but no alterations in mitoATP production, respiration or proton leak compared to *APOE-KO* and *APOE2* indicating that more research is needed to explain all *APOE* genotype dependent alterations in detail.

In contrast to many studies focusing on the comparison of only two *APOE* genotypes, we used the whole set of APOE isogenic lines and included *APOE-KO* in addition to *APOE2*, *APOE3* and *APOE4*. When comparing *APOE-KO* cells to the other *APOE* cell lines, we observed that *APOE-KO* cells have a similar phenotype as *APOE2* but not as *APOE4*, indicating that *APOE4* displays a gain-of-function rather than a loss-of-function effect on astrocyte metabolism. This is in agreement with our previous studies, where both *APOE-KO* iAstrocytes and iN cells showed a similar phenotype to *APOE2* but not to *APOE4*, e.g. in glutamate and Aβ uptake, cholesterol metabolism or inflammatory signaling (de Leeuw et al., 2022; Budny et al., 2024). This is also consistent with the study of Chemparathy and colleagues showing the protective function of *APOE* loss-of-variants in healthy individuals and AD patients (Chemparathy et al., 2024). Furthermore, in our metabolomic enrichment analysis, more glucose- and energy metabolism-related pathways were perturbed in *APOE4* compared to *APOE2* and *APOE-KO* (10 and 11, respectively) than compared to *APOE3* (7). This aligns with other findings, including proteomic data and ATP production, showing larger differences between *APOE4* and *APOE2/APOE-KO* than between *APOE4* and *APOE3*.

## 5. Conclusions

Our study provides compelling evidence that *APOE* variants differentially affect human astrocyte energy metabolism at the proteomic, metabolomic and functional level. Specifically, we demonstrate that *APOE4* drives metabolic dysregulation in iAstrocytes through a gain-of-function mechanism. Despite exhibiting increased mitochondrial respiration and glycolysis, *APOE4* iAstrocytes display reduced ATP production, likely due to mitochondrial dysfunction, as indicated by a significant proton leak. These findings enhance our understanding of AD pathophysiology and highlight potential therapeutic strategies targeting *APOE4*-driven mitochondrial dysfunction in AD.

## Acknowledgments

We appreciate the support of Dr. Alaa Othman and Dr. Martina Zanella from the Functional Genomics Center Zurich in metabolomics analysis. We further thank Maik Krüger für help with Fiji-based nucleus counting.

## Author contribution

**VB:** Conceptualization, Data curation, Formal analysis, Investigation, Methodology, Validation, Visualization, Writing – original draft, Writing – review and editing. **CB:** Investigation. **KJZ:** Investigation. **SMdL:** Methodology, Resources. **RZW:** Data curation, Formal analysis, Visualization. **LR:** Formal analysis, Methodology, Software. **IR:** Methodology. **LFB:** Methodology. **BW:** Resources. **CT:** Conceptualization, Data curation, Formal analysis, Funding acquisition, Methodology, Project administration, Resources, Supervision, Validation, Visualization, Writing – original draft, Writing – review and editing.

