## Supplementary figures for "APOE genotype-dependent differences in human astrocytic energy metabolism"

### Suppl. figure 1

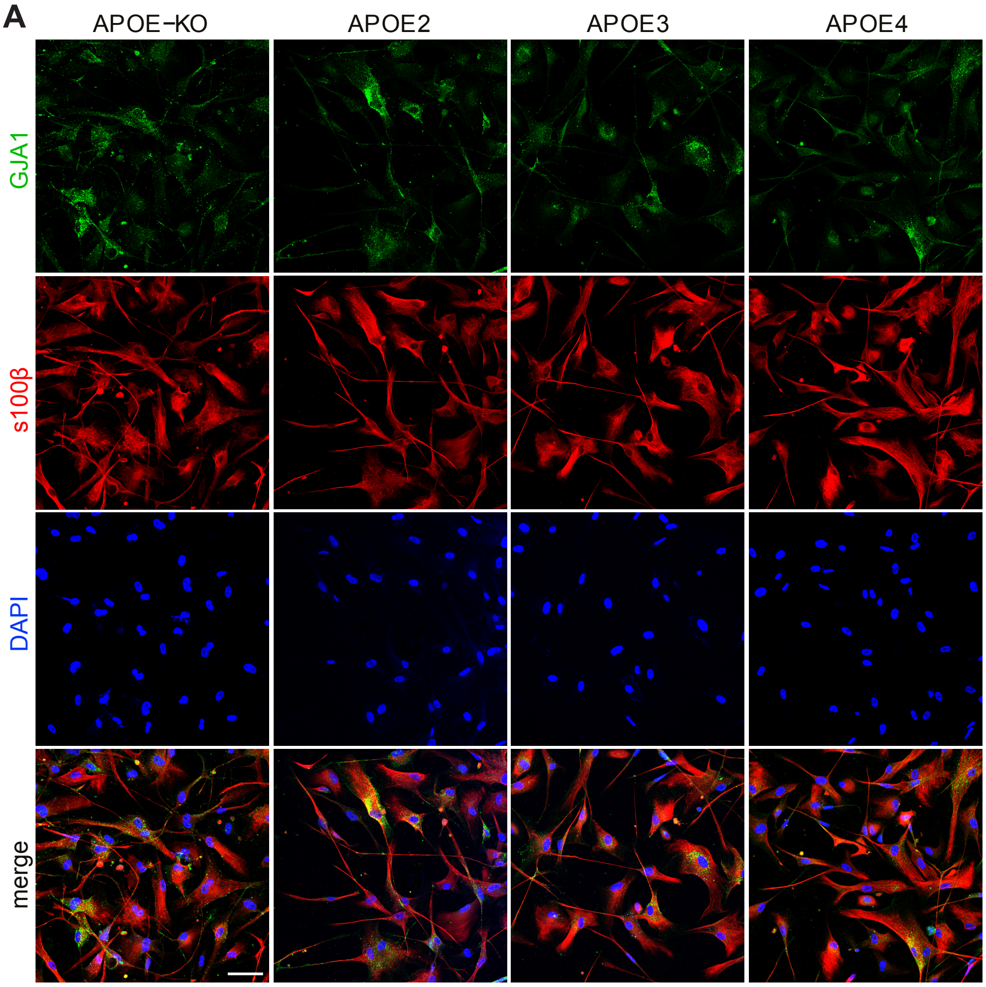


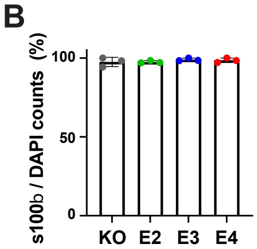


**Supplementary Fig. 1: Characterization of APOE-isogenic iAstrocytes.**
**A:** Confocal images of iAstrocytes at d44, stained for astrocyte markers s100β and GJA1. Scale bar: 100 µm
**B:** Differentiation efficiency plotted as s100β-positive cells per total DAPI count.

### Suppl. figure 2

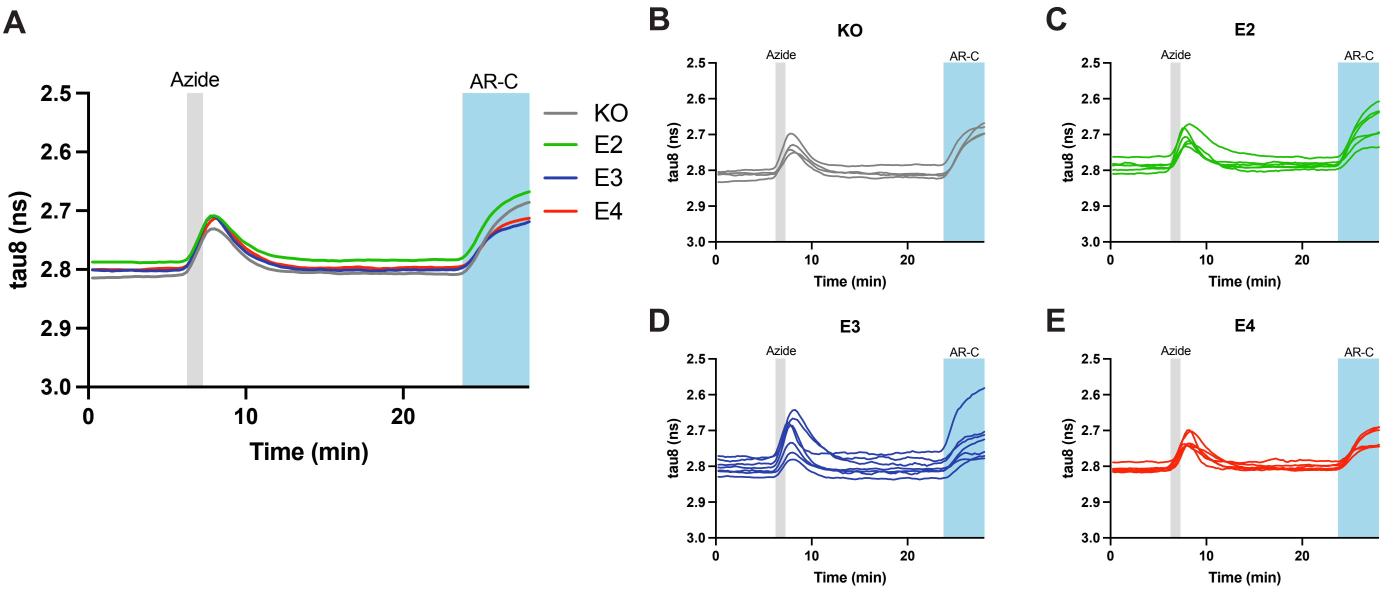


**Supplementary Fig. 2: Complete FLIM measurements (tau8, ns) with the LiLac sensor over time (min).** **A:** Average of all measured lifetimes for each APOE-isogenic cell line over time. **B:** Single lifetime measurements of APOE-KO iAstrocytes over time. **C:** Single lifetime measurements of APOE-E2 iAstrocytes over time. **D:** Single lifetime measurements of APOE-E3 iAstrocytes over time. **E:** Single lifetime measurements of APOE-E4 iAstrocytes over time. Grey=KO, green=E2, blue=E3, red=E4.

### Suppl. figure 3

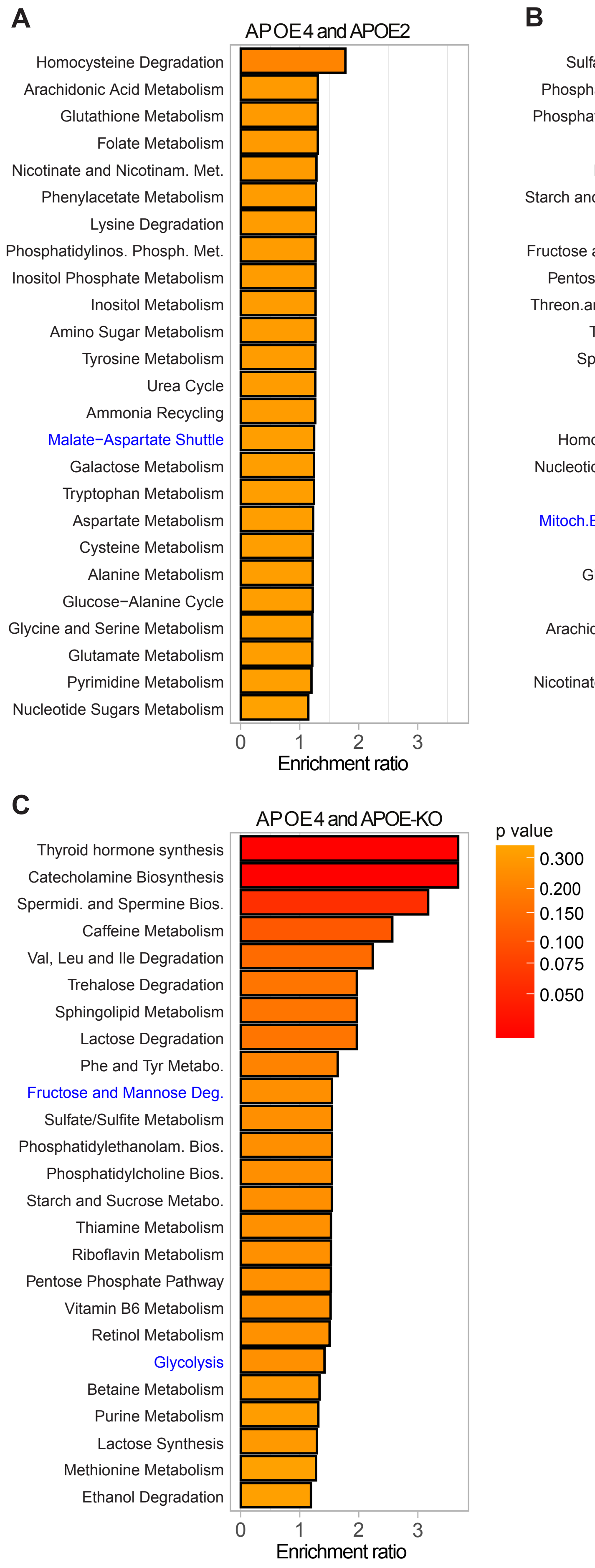


**Supplementary Fig. 3:** Enrichment analysis of metabolic pathways, compared between the respective APOE-isogenic iAstrocytes, analyzed using Metaboanalyst 6.0. Top 25 enriched pathways are shown. Glucose and energy metabolism-related pathwys from Fig 5A-C are highlighted in blue. Pathways are ranked according to the enrichment ratio.

### Suppl. figure 4

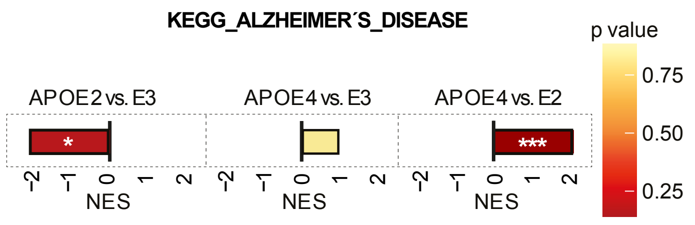


**Supplementary Fig. 4:** GSEA normalized enrichment scores (NES) of Alzheimer's Disease proteomic pathway according to the KEGG database for APOE2 versus APOE3, APOE4 versus APOE3, and APOE4 versus APOE2 iAstrocytes. NES is plotted on the x axis, with color-coded bars for the individual gene ontology (GO) terms. * $p < 0.05$, *** $p < 0.001$
